# Voltammetry in the spleen assesses real-time anti-inflammatory norepinephrine release elicited by autonomic neurostimulation

**DOI:** 10.1101/2022.04.26.489592

**Authors:** Ibrahim T. Mughrabi, Michael Gerber, Naveen Jayaprakash, Santhoshi P. Palandira, Yousef Al-Abed, Timir Datta-Chaudhuri, Corey Smith, Valentin A. Pavlov, Stavros Zanos

**Affiliations:** Institute of Bioelectronic Medicine, The Feinstein Institutes for Medical Research, Manhasset, NY, USA; Zucker School of Medicine at Hofstra/Northwell, Hempstead, NY, USA; Department of Physiology and Biophysics, Case Western Reserve University, Cleveland, OH, USA; Elmezzi Graduate School of Molecular Medicine, Manhasset, NY

**Keywords:** Fast scan cyclic voltammetry, spleen, norepinephrine, inflammation, biomarker, vagus nerve stimulation, splanchnic nerve stimulation, splenic nerve stimulation.

## Abstract

**Background:** The noradrenergic innervation of the spleen is implicated in the autonomic control of inflammation and has been the target of neurostimulation therapies for inflammatory diseases. However, there is no real-time marker of its successful activation, which hinders the optimization of anti-inflammatory neurostimulation therapies and mechanistic studies in anti-inflammatory neural circuits.

**Methods:** In mice, we performed fast-scan cyclic voltammetry (FSCV) in the spleen during intravascular injections of norepinephrine (NE), or during stimulation of the vagus, splanchnic, or splenic nerves. We defined the stimulus-elicited charge generated at the oxidation potential for NE (∼0.8 V) as the “NE voltammetry signal” and quantified the dependence of the signal on NE or nerve stimulation dose. We correlated the NE voltammetry signal in response to splenic nerve stimulation (SpNS) with the latter’s anti-inflammatory effect in a model of lipopolysaccharide- (LPS) induced endotoxemia, quantified as suppression of TNF release.

**Results:** We found that the NE voltammetry signal is proportional to injected amount and estimated peak NE concentration, with 0.3 μM detection threshold. In response to SpNS, the signal increases within seconds, returns to baseline minutes later and is blocked by interventions that deplete NE or inhibit NE release. The signal is elicited by efferent, but not afferent, electrical or optogenetic vagus nerve stimulation, and by splanchnic nerve stimulation. The magnitude of the signal during SpNS is inversely correlated with subsequent TNF suppression in endotoxemia and explains 40% of the variance in TNF measurements.

**Conclusion:** FSCV in the spleen provides a marker for real-time monitoring of anti-inflammatory activation of the splenic innervation during autonomic stimulation.

## Background

The innervation of the spleen by the autonomic nervous system is implicated in the neural control of inflammation [1–3]. Both spinal sympathetic and vagal preganglionic neurons interact with the splenic nerve, and their stimulation results in increased splenic nerve activity and suppression of inflammation [4–12]. Part of this anti-inflammatory effect is initiated by the release of norepinephrine (NE) in the spleen parenchyma by terminals of the splenic nerve, followed by the release of acetylcholine by specialized T-cells and suppression of pro-inflammatory cytokine production by splenic macrophages. These findings prompted the development of new treatments for inflammatory diseases using neurostimulation devices to modulate the activity of autonomic nerves under the umbrella of the growing field of Bioelectronic Medicine [13]. Therapies involving cervical vagus nerve stimulation (VNS) [14], trans-auricular VNS [15, 16], splenic nerve stimulation [8], and ultrasound stimulation of the spleen [5, 6] all rely on the autonomic innervation of the spleen as the common final target to suppress inflammation in patients with inflammatory diseases [5, 6, 9, 11, 12].

As part of the clinical programming of anti-inflammatory bioelectronic therapies, parameters of neurostimulation need to be adjusted on an individual subject basis to maximize anti-inflammatory efficacy, while minimizing off-target effects of neurostimulation [13, 17]. For example, stimulation of the cervical vagus nerve, which also innervates the larynx and the heart, can produce coughing or heart rate changes [18, 19], both undesired responses that could limit anti-inflammatory therapeutic efficacy. To deliver precision anti-inflammatory neuromodulation, the engagement of innervation of end-organs needs to be assessed, ideally in real-time. Whereas markers for the engagement of autonomic innervation of the heart, lungs, and gut have been described [20, 21], no such marker exists for the spleen. A real-time marker for assessing the engagement of the innervation of the spleen to autonomic stimulation could contribute to safer, more effective, and precise anti-inflammatory bioelectronic therapies [13, 17]. Such a marker would also be helpful in mechanistic studies of neural reflexes activated by stimulation of autonomic nerves, something especially useful in rodent studies, in which direct recording of nerve activity evoked by stimulation is challenging because of small nerve size [21, 22].

To address this unmet need, we developed and evaluated a method to assess, in real-time, the release of NE in the spleen in response to different forms of autonomic stimulation. The method is based on fast-scan cyclic voltammetry (FSCV), an electrochemical technique previously used to measure neurotransmitter release in the brain [23, 24] and recently suggested as a potential marker for bioelectronic therapies [25, 26]. We defined a quantitative “NE voltammetry signal” as the total charge generated at the oxidative electrochemical potential for NE (∼0.8 V) in response to a pharmacological or electrical stimulation intervention. We found that the NE voltammetry signal is dose responsive to injected NE and to electrical and optogenetic stimulation of autonomic nerves involved in the innervation of the spleen. The NE voltammetry signal is also responsive to pharmacologic or surgical manipulations that block the effect of neurostimulation on the innervation of the spleen. The magnitude of the NE voltammetry signal significantly predicts the suppression of TNF release by nerve stimulation in acute inflammation, explaining about 40% of the variance in TNF values. Importantly, TNF suppression is greater at low values of the NE signal, and smaller at high values of the NE signal. Our findings indicate that voltammetry of the spleen is a feasible approach to assess, in real-time, the engagement of the splenic innervation by different anti-inflammatory autonomic neurostimulation approaches.

## Methods

### Animals

Male and female C57BL/6 mice, ages 8-16 weeks, were purchased from Charles River Laboratories (Wilmington, MA). ChAT-IRES-Cre (#006410), Vglut2-IRES-Cre (#016963), Ai14 ROSA-tdTomato (#007914) and Ai32 ChR2-eYFP (#024109) mice were purchased from The Jackson Laboratory (Bar Harbor, ME) and crossed to produce ChAT-ChR2-eYFP, Vglut2-ChR2-eYFP, and ChAT-tdTomato mice used for optogenetic and splanchnic nerve stimulation experiments. Animals were housed using a 12- hour light/dark cycle with ad libitum access to food and water. All animal procedures were approved by the Institutional Animal Care and Use Committee (IACUC) of the Feinstein Institutes for Medical Research, NY and complied with relevant NIH policies and guidelines.

### Surgical procedures

Following isoflurane anesthesia, induced at 4% and maintained at 1.5%, a 2-cm midline abdominal incision was made through the linea alba to just above the xyphoid process. The spleen was gently exposed by retracting the stomach and gut to the right while blunt dissecting the gastrosplenic ligament. The lower branch of the splenic neurovascular bundle was identified and gently dissected from surrounding fat. The spleen was then lifted out of the abdominal cavity and supported with cotton tips. A 34-μm diameter carbon fiber microelectrode (CFM; Pinnacle) and an Ag/AgCl reference electrode were mounted on a stereotaxic manipulator and inserted into the exposed spleen. The voltammetry electrodes were advanced ∼0.5 mm into the diaphragmatic surface of the spleen opposite to the inferior branch of the splenic neurovascular bundle supplying the lower half of the spleen (Fig. 1a). To perform splenic nerve stimulation, the splenic neurovascular bundle was placed into a bipolar micro-cuff electrode to deliver electrical stimulation to the splenic nerve, which runs along the splenic artery. To isolate the left cervical vagus nerve, a 1-cm midline incision was made in the anterior neck and the salivary glands were retracted laterally along with the sternocleidomastoid muscle. The vagus nerve was then identified within the carotid sheath, dissected away from connective tissue, and gently placed on a bipolar micro-cuff electrode for electrical stimulation or a custom-made optical cuff for optogenetic stimulation. To perform left splanchnic nerve stimulation, ChAT-tdTomato mice were used to correctly identify the nerve as it enters the celiac-superior mesenteric (CSM) ganglion complex using a fluorescence dissecting microscope. Mice were anesthetized and underwent a midline abdominal incision. The intestines were gently retracted to the right and covered with gauze saturated with warm saline. The celiac artery was then identified, and the left CSM ganglion complex was located around its origin marked by bright red fluorescence. The left splanchnic nerve was traced from the upper left edge of the ganglion back to its thoracic para-vertebral level. The nerve was cuffed with a bipolar electrode (Flex) to deliver electrical stimulation.

**Figure 1.**
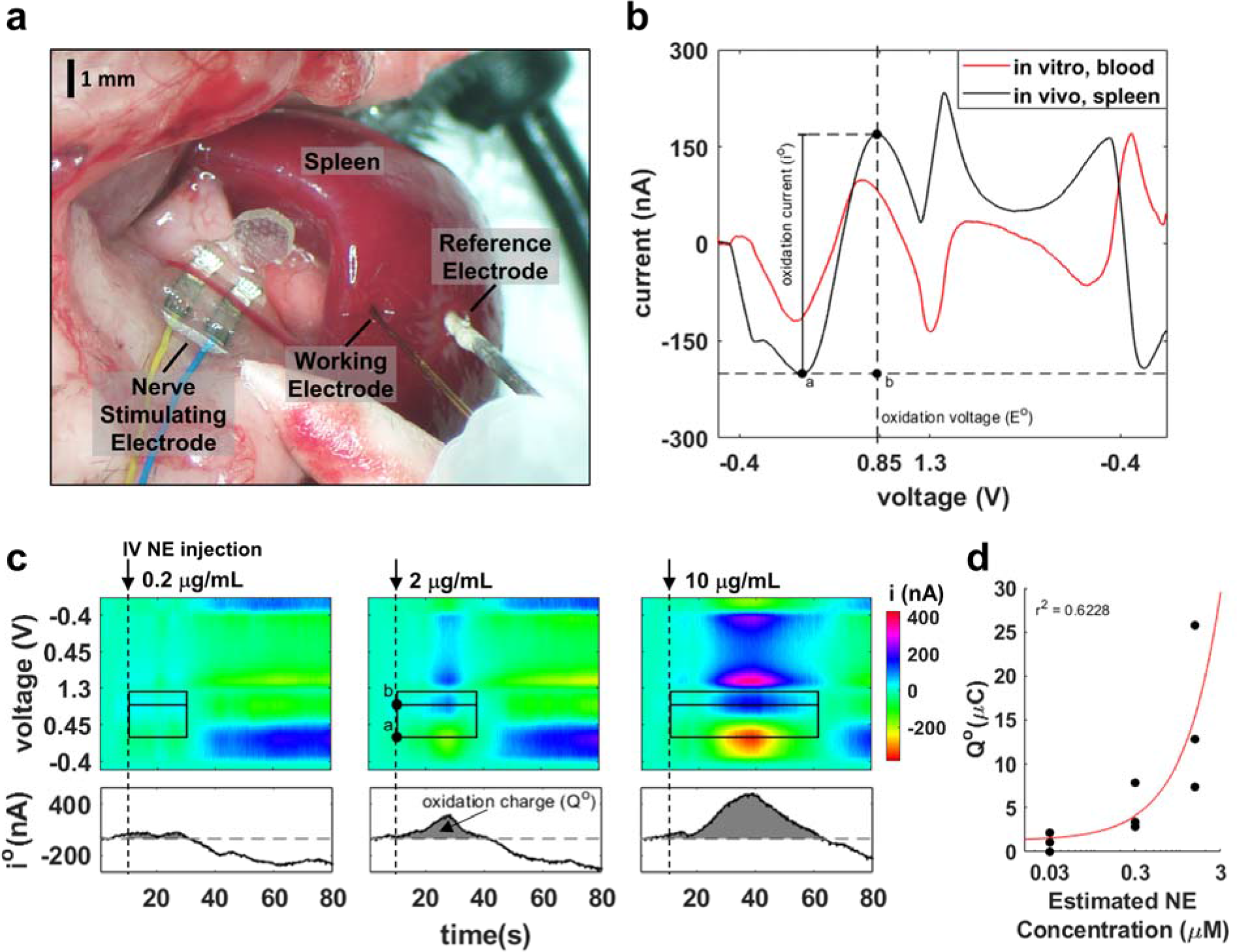
A spleen voltammetry signal responsive to changes in NE. (a) A carbon fiber working electrode and a reference electrode are inserted in the spleen for recording fast scan cyclic voltammetry. A bipolar electrode is placed on the splenic neurovascular bundle for splenic nerve stimulation; in other experiments, the stimulating electrode is placed on the cervical vagus nerve or the splanchnic nerve. (b) Cyclic voltammograms during voltage sweeps (-0.4 V to 1.3 V and back to -0.4 V) in vitro (in heparinized blood, 1 μM NE) and in vivo (in live spleen, 1.5 μM est. peak NE concentration); shown is the NE oxidation voltage (E°) in vivo, recorded at the corresponding peak oxidation current (i°). (c) Representative time-resolved voltammograms during intravenous bolus injection of NE (100 μL). *Top panels*: Current amplitude (represented by color) at different values of sweeping voltage (ordinate), at different times relative to NE injection (abscissa). Outer black box represents the voltage and time boundaries of the peak oxidation current signal (outer boundaries); horizontal line inside box marks the oxidation potential for NE. Dashed vertical line denotes time of drug administration. *Bottom panels*: time-course of oxidation current, calculated as shown in Fig. 1b, during the same time period. Grey-shaded area under the i° trace represents the total oxidation charge (Q°) generated within the time boundaries indicated in the top panels. (d) Q° at multiple NE concentrations measured in 3 animals, at different estimated NE concentrations.

### Fast-scan cyclic voltammetry

A two-electrode commercial system (Pinnacle Technology Inc, Lawrence, KS) was used to perform FSCV using a triangular waveform scanning from -0.4 to 1.3 to -0.4 V at 400 V/sec with a cycling frequency of 10 Hz and -0.4 V holding potential between cycles. Current and voltage data was acquired using the continuous scanning mode at a sampling rate of 1000 Hz. For *in vitro* experiments, blood was collected by cardiac puncture using a heparin-flushed syringe and then pooled together before aliquoting into a 96- well plate. Freshly made NE in PBS solution was added to each well to achieve a final concentration of 0 and 1 μM. After recording background current from the vehicle-treated well, NE was probed in the remaining wells. For *in vivo* experiments, after inserting the voltammetry electrodes into the spleen, the working electrode was conditioned for about 10 minutes before collecting voltammetry data, over which time baseline was typically stable.

### Intravascular infusions

To establish an intravascular (IV) line, the right external jugular vein was isolated and ligated rostrally while blocking blood flow with a loose suture caudally to prevent bleeding. A small incision was then made between the two sutures and a 1-French catheter (Instech Labs, Plymouth Meeting, PA) was advanced into the vessel followed by a 50 μL saline bolus to confirm patency. In some experiments, 100 μL norepinephrine (NE) bitartrate boluses were administered sequentially at escalating doses (10 μg/mL, 2 μg/mL, 0.2 μg/mL) 10 minutes apart. The final concentration of circulating NE after each dose was estimated for an average mouse weight of 25 g based on a volume of distribution of 2 mL estimated to be total blood volume in a mouse [27]. All IV injections were prepared in saline and administered as 100 μL boluses over 10 seconds.

### Nerve stimulation

Electrical stimulation was delivered using commercial bipolar micro-cuff electrodes (CorTec, Freiburg, Germany) or custom-made flexible electrodes (Flex) [28] connected to a stimulus generator (STG4008, Multichannel Systems, Reutlingen, BW Germany). Stimulus trains consisted of bi-phasic charge-balanced square pulses of 10 sec duration, 500 μs pulse width (PW), and 10 Hz frequency at varying intensities. In some experiments, the splenic nerve was blocked by applying a piece of gauze saturated with 2% lidocaine on the splenic neurovascular bundle for 10 min. In cervical vagotomy experiments, the vagus nerve was mechanically stabilized on the micro-cuff electrode by applying a drop of 2-component silicone (Kwik-Sil, World Precision Instruments, Sarasota, FL) and then sectioned either caudal or rostral to the cuff. For optogenetic stimulation experiments, a custom-made optical cuff, consisting of a blue LED light (XLAMP XQ-E, Cree LED, Durham, NC) integrated into a molded silicone cuff, was connected to a stimulus generator (MCS) operating in voltage mode. Optical stimulation was delivered using 10-sec stimulus trains of 10 ms PW and 30 Hz frequency while varying intensity by changing the voltage driving the LED. During vagus stimulation experiments, animals were instrumented with ECG leads and nasal airflow temperature sensors to monitor heart rate and breathing rate responses.

### Norepinephrine depletion

Reserpine in DMSO (Tocris Bioscience, Bristol, UK) was diluted in saline and administered to animals by i.p. injection (5 mg/kg). 18-24 hours post-injection, NE elicited by SpNS was probed using voltammetry in the spleen as described.

### Endotoxemia model

Following isoflurane anesthesia, the spleen was exposed through a 1-cm lateral abdominal incision just below the left costal margin and instrumented with voltammetry electrodes as described before. After 10 min of CFM conditioning, the splenic neurovascular bundle was electrically stimulated for 3 min (300-500 μA, 500 μs, 10 Hz) using a bi-polar cuff electrode (CorTec) while performing spleen FSCV. To ensure complete recording of the elicited NE signal, FSCV sweeping was continued for at least 10 min post-stimulation and the disappearance of the signal was verified before stopping the recording. The voltammetry electrodes were then carefully removed, and the micro-puncture site was repaired with tissue glue. The abdominal incision was closed in two layers, and the animal was allowed to recover from anesthesia. Two hours post-stimulation, animals were injected intraperitoneally with 1 mg/kg of lipopolysaccharide (LPS endotoxin from *E. coli* 0111:B4 ultra-pure; InvivoGen, San Diego, CA) dissolved in saline. Heparinized blood was collected by cardiac puncture under terminal anesthesia 90 min post-LPS injection and centrifuged immediately at 2000 *x g* for 10 min. Plasma TNF-α was determined using TNF-α ELISA kit (Invitrogen, Waltham, MA) following manufacturer’s instructions.

### Data analysis and statistics

Voltammograms were background-subtracted and smoothed with a moving average filter. Oxidation current (i°) was determined in each voltammetry cycle as the difference between the trough and the peak of the current trace during the oxidation part of the cycle (points a and b, respectively, Fig. 1b). The “NE voltammetry signal” was defined as the integral of i° over time during an integration window, corresponding to the total oxidation charge (Q°) (Fig. 1c). This method was chosen because peak and duration of the NE transient in response to a stimulus are both likely to be important for downstream biological effects of NE release. The i° integration window was set between the stimulus trigger and a second time boundary, which was determined by an algorithm that calculates the time at which the i° trace returns to baseline. The algorithm for determining the window for integration of i° was implemented in MATLAB (MathWorks, Natick, MA) and is described in detail in the Supplementary Methods.

For statistics, paired, two-tailed t-test was used to determine significance in comparisons between two groups in experiments where treatment and control was done within the same animal (i.e. lidocaine) and unpaired t-test was used for comparisons between animals (i.e. reserpine, LPS). MATLAB’s model fitting package was used for linear regression analysis and computation of r^2^ for stimulation intensity vs Q° in splenic stim and VNS experiments, and in Q° vs TNF a power model was used for regression instead due to the asymptote at Q° = 0 causing non-linearity. Correlation between Q° and TNF was tested with Spearman’s ranked correlation coefficient due to the aforementioned non-linearity. P values less than 0.05 were considered significant. All calculations were done in MATLAB.

## Results

### A voltammetry signal in the spleen represents norepinephrine release

To define a physiological marker that assesses the release of norepinephrine (NE) in the spleen, we first performed fast-scan cyclic voltammetry (FSCV) in vitro to establish the oxidation potential of NE in blood. We found that the oxidation potential of NE (E°), the potential at which peak oxidation current occurs, is 0.71 V in blood (at 1 μM NE concentration; Fig. 1b), in agreement with prior reports [25]. To determine the oxidation potential E° in the spleen and assess whether oxidation current (i°) tracks transient changes of NE concentration, we performed FSCV in the spleen during intravenous injections of NE amounts estimated to produce peak concentrations of 0.03-1.5 μM in circulating blood. During NE infusion, i° rises and then returns to baseline, as opposed to saline infusion where i° remains unchanged (Fig. S1a-c); E° during NE infusion is 0.85 ± 0.06 V (n = 5 experiments) (Fig. 1b). Oxidation current (i°) starts increasing at approximately 0.2 μg/mL NE injection and increases further with increasing injected amounts of NE (Fig. 1c). To capture both the magnitude and the duration of the transient change in NE, we defined the “NE voltammetry signal” as the total oxidation charge (Q°), calculated as the area under the i° trace, between the time of NE injection and the time at which i° drops to pre-injection baseline level (Fig. 1c, black boxes). Q° is linearly proportional to the injected amount of NE (Fig. S1d) and the estimated peak NE concentration in blood (semi-log plot in Fig. 1d). From this data, we estimate the detection threshold of Q° for NE to be 0.3 μM NE, or less (Fig. 1d).

Together, these findings indicate that NE can be detected in vivo by voltammetry in the spleen, and that the NE voltammetry signal is proportional to the injected amount and the resulting (estimated) concentration of NE.

### NE voltammetry signal is responsive to bioelectronic activation of the splenic nerve

Electrical stimulation of the splenic nerve (SpNS) elicits release of NE in the spleen [8]. To test whether the spleen voltammetry signal tracks NE release under these conditions, we performed FSCV in the spleen during SpNS (Fig. 1a). During SpNS, i° rises and then settles back to baseline, as opposed to sham stimulation where i° remains unchanged (Fig. S2a-c). The shape of the voltammogram is similar to that recorded during the NE infusion experiments. The magnitude and duration of the i° trace both increase with increasing intensity of the electrical stimulus (Fig. 2a); expectedly, Q° also increases with stimulus intensity (Fig. 2b).

**Figure 2.**
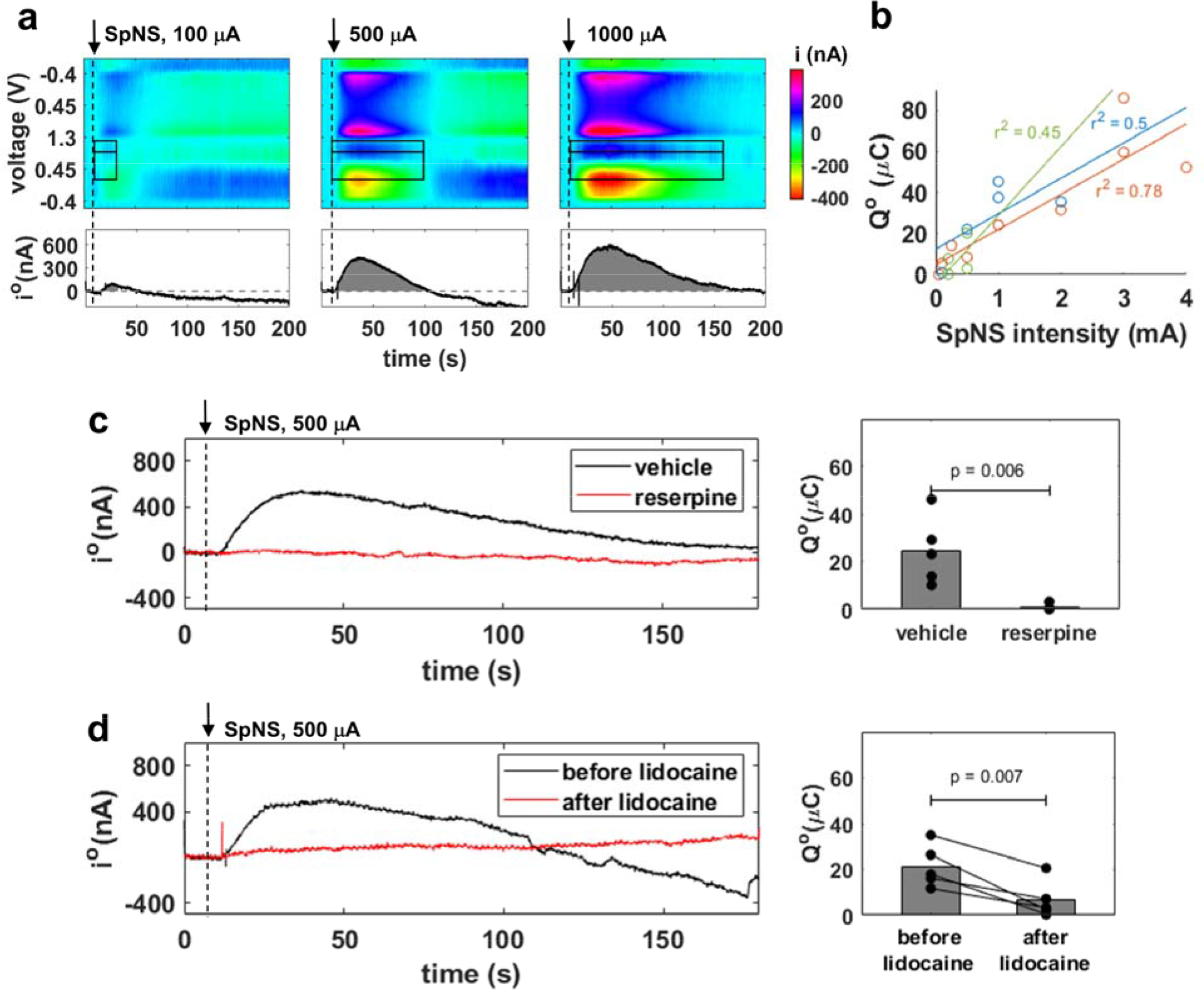
Activation of the splenic nerve by splenic nerve stimulation (SpNS) elicits a NE voltammetry signal in the spleen. (a) NE voltammetry signal elicited by SpNS (10 sec duration, 500 μs pulse width, 10 Hz pulsing frequency). *Top and bottom panels* similar to those of Fig. 1c. Dashed vertical line indicates the onset of SpNS. (b) Oxidation charge (Q°) vs. stimulation intensity in 3 animals, each represented by a different color; lines represent least square fits on data from each animal and r^2^ is the respective coefficients of determination. (c) *Left panel:* Time course of NE peak oxidation current (i°) in response to SpNS, in reserpine- and sham-treated mice. *Right panel:* Q° in response to SpNS in reserpine-treated vs. vehicle-treated animals (n = 4 in each group). (d) *Left panel:* Time course of i° in response to SpNS recorded in the same animal before and after the topical application of lidocaine on the splenic nerve. *Right panel:* c before and after lidocaine application, in 4 animals.

To verify that the voltammetry signal elicited from SpNS is, at least partly, mediated by NE release, we performed FSCV during SpNS in animals treated with reserpine to deplete monoamines [4]. The NE voltammetry signal is absent in reserpine-treated animals; in contrast, it is evoked in vehicle-treated animals (Fig. 2c). To verify that the source of released NE is indeed the stimulated splenic nerve, SpNS was delivered before and after lidocaine, a local anesthetic agent blocking nerve depolarization, was directly applied on the splenic nerve. The NE voltammetry signal after lidocaine application is significantly smaller compared to the signal before lidocaine (Fig. 2d).

Together, these findings indicate that the NE voltammetry signal in the spleen represents the release of NE in response to activation of the splenic nerve by SpNS.

Dashed vertical line indicates the onset of SpNS.

### NE voltammetry signal is responsive to bioelectronic activation of the pre- ganglionic efferent vagus and splanchnic nerves

Vagus nerve stimulation (VNS) elicits norepinephrine (NE) release in the spleen via the efferent arc of the inflammatory reflex [29]. To determine whether the NE voltammetry signal is responsive to VNS, we performed FSCV in the spleen during cervical VNS. During VNS, i° rises and then settles back to baseline, as opposed to sham stimulation where i° remains unchanged (Fig. 3a & S3a-c); the magnitude of i°, and of its derivative measure Q°, are dose-responsive to VNS intensity (Fig. 3a & b). Notably, the voltammetry signal in response to VNS has lower amplitude and shorter duration compared to that elicited by SpNS (Fig. S2a-c & S3a-c).

**Figure 3.**
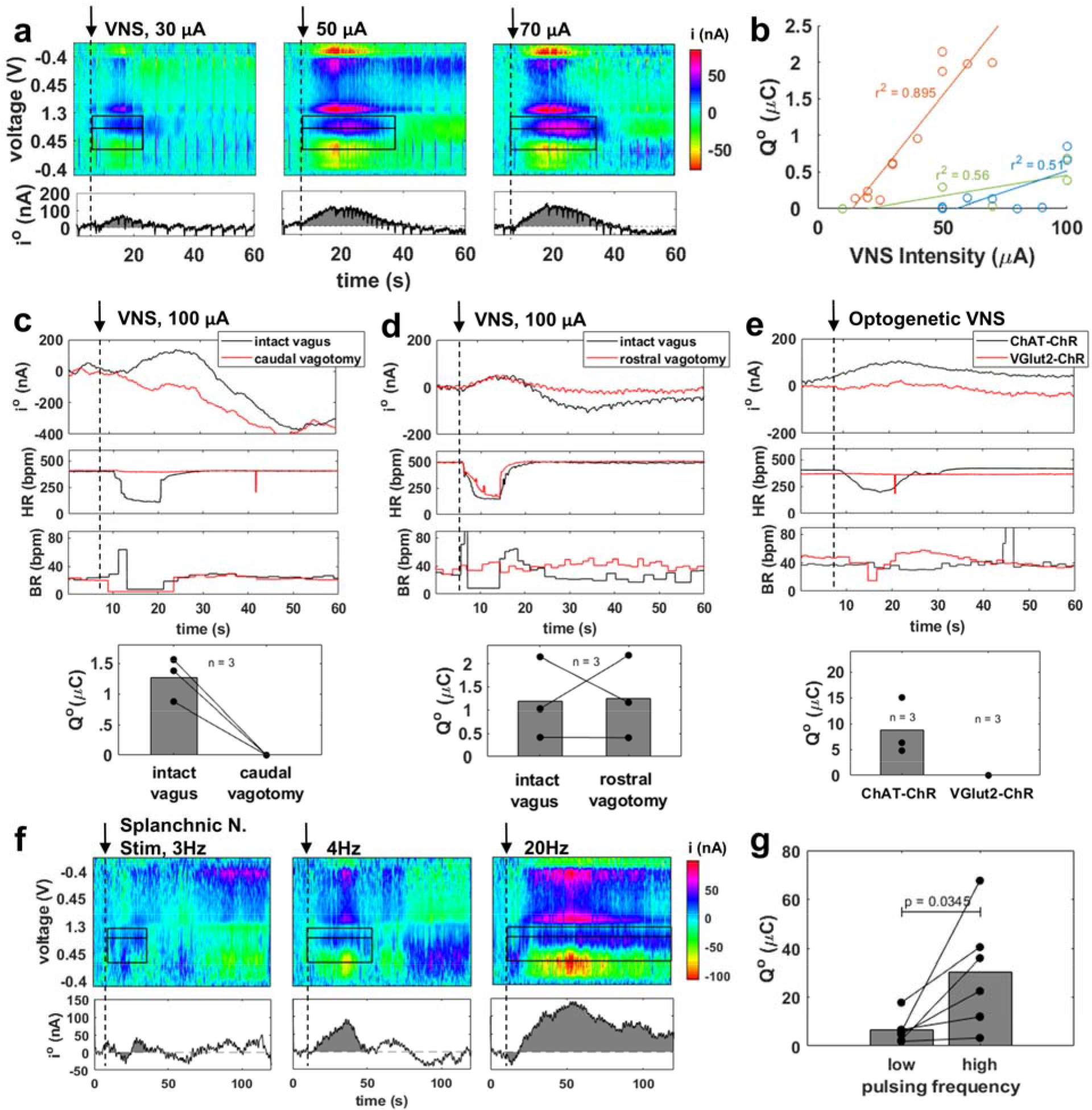
Stimulation of the efferent, but not afferent, vagus or of the splanchnic nerve elicits a NE voltammetry signal in the spleen. (a) Representative time-resolved voltammograms elicited by electrical left vagus nerve stimulation (VNS) (10-100 μA intensity, 500 μs pulse width, 10 Hz frequency, 10 sec duration). *Top and bottom panels* are similar to those in Fig. 1c. The dashed vertical line denotes onset of VNS. (b) NE oxidation charge (Q°) in response to different VNS intensities in 3 animals; lines represent least square fits on data from each animal and r2 the respective coefficients of determination. (c) Time course of NE oxidation current (i°) elicited by VNS (top panel), associated heart rate and breathing rate responses (2 middle panels); Q° values are shown in bottom panel, in 3 animals before and after caudal vagotomy. (d) Same as (c), but for rostral vagotomy. (e) Representative NE oxidation current trace elicited by optogenetic VNS in ChAT-ChR or in Vglut2-ChR mice (top panel), along with the associated heart rate and breathing rate responses (2 middle panels); Q° values from 3 animals in each case are shown in bottom panel. (f) Representative time-resolved voltammograms elicited by electrical left splanchnic nerve stimulation (500 μA intensity, 100 μs pulse width, 2-20 Hz frequency, 10 sec duration). *Top and bottom panels* are similar to those in Fig. 1c. (g) Q° values during splanchnic nerve stimulation using low (2-4 Hz) and high (20 Hz) pulsing frequency in the same animal (n = 6).

To determine whether activation of afferent (sensory) or efferent (motor) vagus nerve is responsible for the observed NE voltammetry signal, VNS was delivered before and after dissection of the vagus nerve, above or below the level of stimulation (caudal or rostral vagotomy, respectively). In response to cervical VNS with the vagus nerve intact, activation of afferent vagal fibers elicits changes in breathing and of efferent vagal fibers in heart rate [30]. When VNS is delivered after caudal vagotomy, the heart rate response disappears, and the NE voltammetry signal is diminished (Fig. 3c). In contrast, when VNS is delivered after rostral vagotomy, the breathing response disappears but the NE voltammetry signal is not affected (Fig. 3d). Likewise, selective activation of efferent vagal fibers by optogenetic VNS in ChAT-ChR mice, elicits the NE signal in the voltammogram. In contrast, selective activation of afferent vagal fibers, by optogenetic VNS in Vglut2-ChR mice, does not elicit a NE signal. (Fig. 3e).

Splanchnic nerves provide sympathetic pre-ganglionic fibers to celiac-superior mesenteric ganglion neurons whose axons form the splenic nerve releasing NE to the spleen [31–34]. To determine whether activation of these preganglionic fibers results in a NE voltammetry signal in the spleen, we used fluorescence imaging in ChAT-tdTomato mice to isolate the splanchnic nerves (Fig. S3d) and delivered electrical splanchnic nerve stimulation using a cuff electrode. We found that stimulation of the splanchnic nerves elicits the NE voltammetry signal in the spleen (Fig. 3f). Splanchnic nerve stimulation at high pulsing frequency (20 Hz) consistently produces higher Q° than at low pulsing frequency (2-4 Hz), indicating that the voltammetry signal is dose-responsive (Fig. 3g).

### NE voltammetry signal during splenic nerve stimulation is predictive of subsequent TNF suppression in LPS endotoxemia

To determine whether the NE voltammetry signal in the spleen can assess the engagement of the anti-inflammatory neuroimmune pathway, Q° was measured during splenic nerve stimulation (SpNS), followed by injection of LPS and, 90 minutes later, measurement of TNF from plasma samples (Fig. 4a). As expected from previous studies [8, 9, 12], we found that, overall, SpNS elicits suppression of TNF compared to sham stimulation (Fig. 4b). Importantly, we found that Q° is inversely correlated with the degree of TNF suppression: smaller values of Q° are associated with TNF suppression, whereas greater values of Q° are associated with TNF values comparable to sham stimulation. As a result, about 40% of the variance in TNF values is explained by Q° (Fig. 4c).

**Figure 4.**
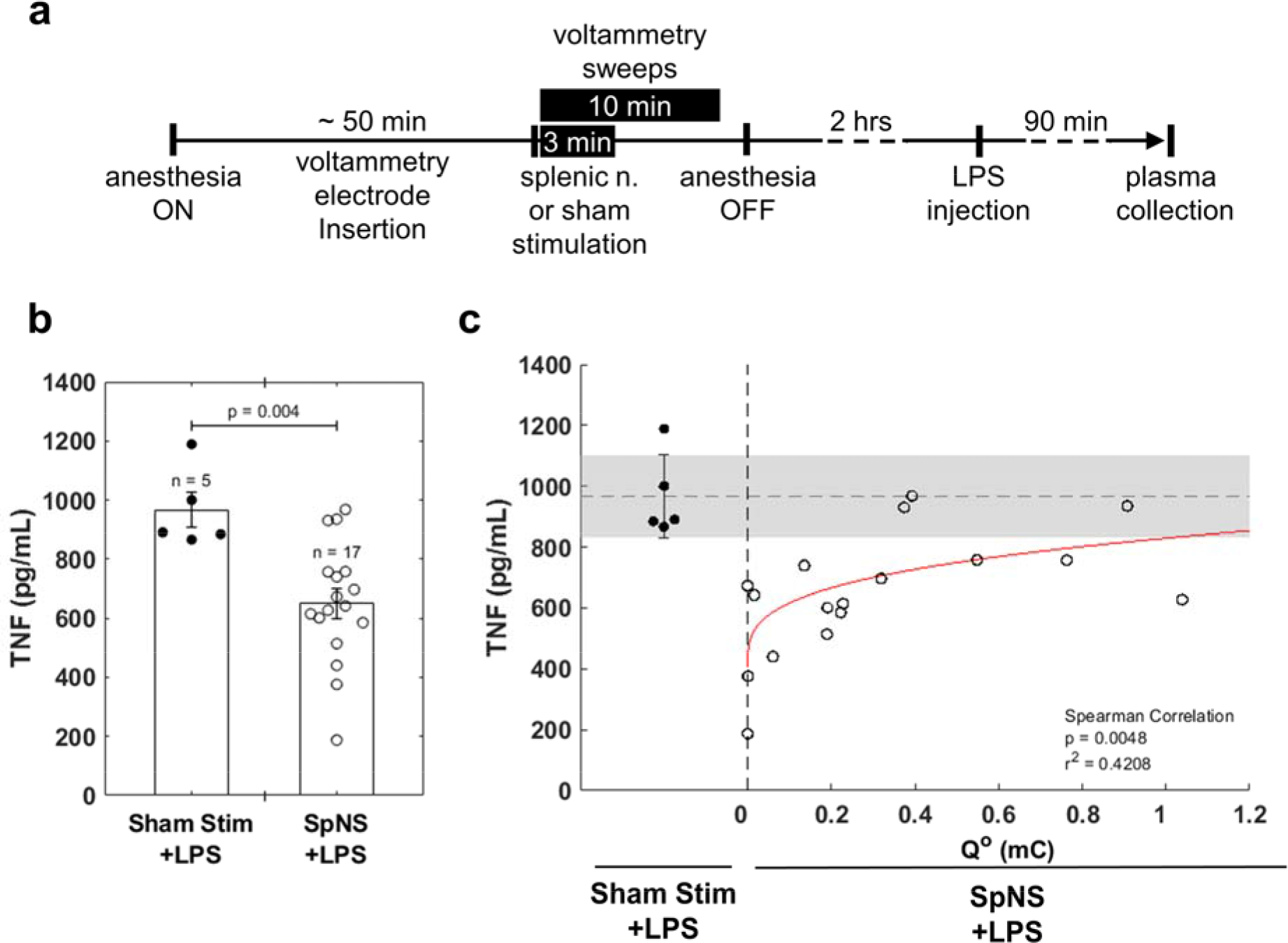
NE voltammetry signal during splenic nerve stimulation (SpNS) is predictive of subsequent TNF suppression in LPS endotoxemia. (a) Schematic representation of experimental timeline. Mice were anesthetized and instrumented with voltammetry electrodes. SpNS was delivered for 3 min (300-500 μA, 500 μs, 10Hz) while performing voltammetry for 10 min to capture full signal. Lipopolysaccharide (LPS) was injected 2 hours after SpNS and blood was collected 90 min post-LPS injection. (b) Plasma TNF levels 90 min post-LPS injection from animals that received SpNS compared with sham stimulated animals. (c) Q° values, measured during SpNS vs. levels of TNF in plasma, measured 90 min after LPS injection, in the same animals. In sham-stimulated animals, Q° is not reported, as it is not defined in the absence of a stimulus.

## Discussion

Autonomic neurostimulation therapies aim to modulate neural circuits controlling inflammation, of which a significant component is the innervation of the spleen. Assessing the preclinical effectiveness of these therapies has relied on indirect measurements, i.e., levels of TNF after an LPS challenge, which are typically made hours after the stimulus has been delivered, oftentimes in terminal experiments [35]; such measurements cannot be used to assess engagement of the anti-inflammatory pathway in real-time. Our work indicates that voltammetry in the spleen is an inexpensive and reproducible method to directly assess engagement of the anti-inflammatory pathway in response to autonomic stimulation. It also demonstrates for the first time a quantitative relationship between a physiological biomarker that is registered in real-time, the NE voltammetry signal, and an immune-mediated response that is several steps downstream, namely TNF suppression in endotoxemia. Physiological markers of anti-inflammatory effectiveness could help in the implementation of precision autonomic neuromodulation therapies, by allowing calibration of stimulation parameters to maximize desired and minimize undesired, off-target, effects in individual subjects [17].

Norepinephrine (NE) is the main neurotransmitter of the sympathetic nervous system, and the neurotransmitter released by fibers of the splenic nerve [36]. It belongs to a group of electrically active compounds termed catecholamines, which include epinephrine and dopamine [37]. Due to its oxidation-reduction properties, changes in NE concentration can be detected almost in real-time by electrochemical methods, including cyclic voltammetry. Fast scan cyclic voltammetry (FSCV) can detect local changes in NE concentration almost in real time owing to its high cycling speed (400 V/s at 10 Hz), and at low concentrations with increased spatial selectivity due to the microscopic scale of the working electrode [38]. When measuring in vitro in blood, we observed an E° of approximately ∼0.7 V, which agrees with previous reports [25]. When measuring from the spleen during intravenous NE injections, E° is on average 0.85V +/- 0.06 (Fig. 1b). One possible reason for the increase of E° in tissue and blood compared to PBS (0.6 V), is biofouling, a common phenomenon when electrochemical probes are implanted in biological tissues [39].

During intravenous injection of NE, i° rises within seconds and returns to baseline after several minutes. Furthermore, i° and its derivative measure Q° are dose-responsive with regard to the amount of injected NE (Fig. 1c-d). During saline injection or sham stimulation conditions i° either did not change or it slowly drifted monotonically akin to a baseline drift (Fig. S1c & S2c), but never exhibited transients changes seen during stimulation. Using intravenous injections of different amounts of NE, we found that 0.03 μM is the lowest (estimated) concentration that produces a voltammetry signal and at 0.3 μM a robust signal is always produced, indicating that detection threshold of our method is between 0.03-0.3 μM. To estimate the NE concentration in blood for a given amount of injected NE, we used the average blood volume of a mouse as the volume of distribution of NE; however, we did not directly measure the concentration of NE, this being a limitation of our study. Compared to previous reports, in which 0.03-0.06 μM of NE were detected after neurostimulation [8, 9, 40, 41], the detection threshold of our system is relatively high.

However, NE concentrations in previous reports were determined in processed blood samples using ELISA or mass spectrometry, where sample collection and processing to collect plasma or serum is typically done 5-10 minutes after stimulation. For that reason, NE concentrations at the time we perform voltammetry is likely greater, and above the detection threshold of the voltammetry method. Further, the concentration of NE in the spleen parenchyma is likely higher than plasma levels, since only a small portion of it overflows to the circulation [42].

The motor arc of a well-studied neuroimmune pathway that controls inflammation starts with parasympathetic efferent vagal fibers and sympathetic efferent splanchnic nerve fibers synapsing on noradrenergic neurons of abdominal ganglia and continues with post-ganglionic noradrenergic fibers inside the splenic nerve [1, 3, 43–45]. Activation of this neuroimmune pathway via neurostimulation of the vagus, the splanchnic or the splenic nerve and its terminals results in activation of splenic nerve fibers, release of NE in the spleen and subsequent suppression of manifestations of acute inflammatory responses [6, 8, 46–48]. Release of NE from splenic nerve fibers has been measured using biochemical assays measuring NE in splenic homogenates or inferred by recording splenic nerve activity after neurostimulation [4, 29]. However, such techniques cannot assess the real-time release dynamics of NE. In this study, we used voltammetry in the spleen to record a transient NE signal that is responsive to autonomic neurostimulation of several nerve targets that activate the anti-inflammatory neuroimmune pathway and suppress inflammation. For that reason, FSCV may be used in mechanistic studies of the regulation of inflammation by different components of the autonomic nervous system.

The NE voltammetry signal is responsive to stimulation of the splenic nerve (SpNS). The signal is proportional to stimulus intensity (Fig. 2a) and is suppressed by manipulations that block nerve activation by stimuli (Fig. 2d) or release of NE upon nerve activation (Fig. 2c). The NE signal appears within seconds after the onset of SpNS and lasts from several seconds to several minutes, depending on stimulus intensity (Fig. 2a). This is consistent with previous studies in pigs and in human post-mortem specimens, in which splenic nerve evoked compound action potentials were found to increase with higher stimulation intensity or duration and were blocked by lidocaine [8, 9]. The amount of NE released into splenic venous outflow during SpNS was shown to increase with higher stimulation charge [9].

Furthermore, those studies showed that physiologic responses to SpNS, such as changes in blood pressure, last for more than a minute after stimulation [8, 9], resembling the kinetics of the elicited NE signal in our study. The NE signal in our study has a slower and longer time course than what is reported in voltammetry studies in the brain (e.g. [23, 49–51]). In the brain, due to the high density of neural processes, the voltammetry electrode has an intimate contact with nerve terminals and NE is almost immediately detected. In contrast, in the spleen, NE likely has to travel through the red pulp cords, where nerve fibers form a network [44], before sufficient concentration builds up at the voltammetry electrode. The majority of released NE in the spleen is cleared by reuptake via the high-affinity, low-capacity NE transporter (NET) [42, 52]. NET is inhibited by electrical stimulation [53], which could explain the relatively long duration of the NE voltammetry signal following SpNS.

We also found that the NE voltammetry signal is responsive to stimulation of the vagus nerve (Fig. 3a). This is consistent with previous observations that VNS exerts anti-inflammatory actions via a splenic nerve-dependent mechanism [4]. In addition, we found that the voltammetry signal is responsive to efferent but not afferent VNS (Fig. 3c-e). This is consistent with previous reports implicating efferent vagal fibers in relaying signals to the splenic nerve via the celiac-superior mesenteric ganglion complex [3, 7]. It is likely that afferent vagal fibers may also contribute to the anti-inflammatory effect, via delayed engagement of multi-synaptic, vagal-sympathetic and vagal-vagal reflexes [32, 33, 54]; however, the lack of a temporally precise volley of action potentials reaching the spleen in response to afferent VNS may be responsible for the lack of a clear NE voltammetry signal. The voltammetry signal after efferent VNS has consistently lower magnitude than that after splenic nerve stimulation, even after stimulation at high intensities that produce a significant drop in heart rate (Fig. 2b-3b). A possible explanation may be that VNS results in eliciting action potentials only on a subset of postganglionic splenic nerve fibers or in activating those fibers at a sub-maximal degree. This implies that VNS at relatively low intensities might elicit release of NE at levels below the limit of detection of the voltammetry method. Studies that demonstrated anti-inflammatory actions of VNS at levels below those that induce a reduction in heart rate support this notion [47].

Third, the NE voltammetry signal is responsive to direct stimulation of the splanchnic nerve (Fig. 3f). To demonstrate this, we used a ChAT-tdTomato mouse strain to visualize and isolate the celiac-superior mesenteric ganglion complex and the associated splanchnic nerve. Although this ganglion and the splanchnic nerve have been isolated under direct vision [33, 55], we report here, for the first time, that the use of fluorescence microscopy improves yield and accuracy. The finding that splanchnic nerve stimulation elicits a NE signal in the spleen is consistent with reports that implicate splanchnic nerve activity in the anti-inflammatory effect of VNS [31–34]. For example, administering VNS with the splanchnic nerve sectioned abolishes the TNF-lowering effect of stimulation in a model of LPS endotoxemia [33].

Notably, the magnitude of the NE voltammetry signal varies across animals, regardless of the stimulated nerve. This variability may arise because of differences in electrode placement directly affecting the number of activated nerve fibers, even at identical stimulus intensities. This underlines the need for using quantifiable markers of target engagement for calibration of the dose of bioelectronic therapies [17, 21]. Another source of signal variability may lie with the voltammetry technique itself. It is likely that the location of the voltammetry electrode in the spleen relative to nerve endings influences the signal. The noradrenergic innervation of the spleen has a mesh-like structure and is most dense towards the center of the organ [43, 44]. Although the working length of the voltammetry electrode was the same in all of our experiments (500 μm), the exact insertion depth into the spleen likely varied, affecting which anatomical compartment of the spleen was sampled. This source of variability is a limitation of single-electrode cyclic voltammetry; it will have to be quantified in future studies with multiple voltammetry electrodes, capturing NE voltammetry signals from several sites in the spleen.

Previous studies of LPS endotoxemia report considerable variability in TNF inhibition in response to VNS or SpNS [12, 35, 54]. Accordingly, we found that the same intensity of SpNS produces a wide range of TNF responses in animals injected with LPS (Fig.4b); out of 17 stimulated animals, at least 3 have TNF values similar to those of sham-stimulated animals. The magnitude of the NE voltammetry signal during SpNS explains a significant portion of this variability, about 40%. In addition, TNF suppression is more likely to occur when stimulation results in relatively small Q° values but is often minimal with greater values of Q° (Fig. 4). This relationship has not been previously described, and we can only speculate of the mechanism behind it. Monocytes/macrophages are known to express both α- and β-adrenergic receptors (ARs); α-ARs are typically associated with mediating pro-inflammatory signaling and bind NE with high affinity at low concentrations, whereas β-ARs mediate anti-inflammatory effects and only bind NE at high concentrations [56–59]. Upon SpNS, NE concentration is highest at nerve terminals, where it is released, and lower away from nerve terminals [57]. Therefore, the location of immune cells relative to nerve terminals may affect the anti-inflammatory response to SpNS. It is possible to consider that relatively small NE release may only affect ChAT^+^ T cells that lie close to nerve terminals by binding β_2_- ARs and causing acetylcholine release, which then acts on macrophages to inhibit TNF release. In contrast, greater NE release may result in monocytes/macrophages located further away, in the red pulp of the spleen, to be exposed to small but not zero concentrations of NE, binding high-affinity α-ARs and favoring TNF release [44, 57]. In fact, in a similar model of endotoxemia, one study found that co-treating animals with both α- and β -AR agonists resulted in TNF suppression similar to that in vehicle treated animals, while treating with an α- or β-AR agonist alone increased or decreased TNF, respectively [59].

Furthermore, large stimulus intensities or pulsing frequencies might induce the release of co-transmitters, such as NPY and ATP, that are also immune-modulators and might negate the NE effect [57, 60].

Therefore, it is conceivable that non-responders to neurostimulation, reported frequently in previous studies, could be in fact animals that were under- or over-stimulated. In our study, which produced a range of Q° values, animals with high Q° values showed smaller TNF suppression (Fig. 4C). In a study by Brinkman et al. [12], most of the animals receiving splenic nerve stimulation had TNF values close to those of the sham-stimulated group (∼6000 pg/mL vs. ∼7500 pg/mL, respectively). The results from our study, which used similar stimulation parameters, suggest that in the Brinkman et al. study it is likely that in a subset of animals that showed smaller TNF suppression, Q° may have been similarly high. These findings underscore the potential usefulness of FSCV as a tool to calibrate stimulation dose and optimize intensity and other stimulation parameters to achieve a predictable anti-inflammatory response within and across subjects. Although recent studies in pigs suggest using changes in splenic artery flow as a marker of effective splenic nerve stimulation [8], this approach is indirect and does not reflect the actual NE content in the spleen. Changes in splenic artery blood flow could be mediated by the direct effects of stimulation on the vessel innervation itself [61], and not necessarily reflect activation of intrinsic splenic nerve fibers.

The FSCV method has several limitations. First, the method is invasive, as it requires puncture and repair of the spleen, which might alter the splenic response and introduce an additional source of variability.

This variability is likely to be minor, as the range of TNF responses we observed is similar to previously reported data using the same LPS and concentration [47]. Second, our configuration has a relatively high threshold for detection of NE, which may limit its use in some neuromodulation therapies, for example, auricular or low-level VNS [62–64]. Third, due to variations in the voltammetry electrode placement, measurement of Q° has low spatial resolution and differences in signal magnitude may partially represent variations in electrode location relative to intrinsic nerves or splenic blood supply. Finally, FSCV using our methodology cannot distinguish between different catecholamines (e.g. NE and dopamine) due to their similar chemical structures [65]. However, since dopamine is typically co-released with NE in very small amounts [66], its contribution to the signal is likely minimal.

Recent advances in voltammetry electrode fabrication allow integration onto highly flexible biocompatible materials and its implantation into various organs to record catecholamine transients. For example, the use of platinum wires allowed the detection of real-time NE release in a beating heart overcoming the fragile nature of carbon fibers [26]. Further, Li et al. developed a flexible and stretchable multichannel interface integrated on a graphene-elastomer composite, which they used to measure monoamine transients chronically in the brain and gut [67]. These technological advances, along with progress in the processing and analysis of voltammograms, may facilitate the development of implanted devices that continuously monitor catecholamine release in the spleen as part of integrated autonomic stimulation systems [68].

## Conclusion

Tools to monitor the anti-inflammatory effectiveness of neurostimulation are needed to calibrate and adjust neurostimulation dose and develop precision bioelectronic therapies. FSCV in the spleen detects in real-time NE release in response to autonomic neurostimulation, which correlates with engagement of the anti-inflammatory neuroimmune pathway. FSCV can be used pre-clinically to resolve neural circuits activated by existing neurostimulation approaches, to identify new targets for autonomic neuromodulation, and to optimize stimulation parameters with regard to the anti-inflammatory effect. With additional development and validation, FSCV could be used clinically to provide a continuous readout associated with anti-inflammatory therapeutic efficacy.

## Abbreviations

FSCV: Fast scan cyclic voltammetry; NE: Norepinephrine; SpNV: Splenic neurovascular bundle; PBS: Phosphate buffered saline; SpNS: Splenic nerve stimulation; E°: oxidation potential; i°: oxidation current ; Q°: total oxidation charge; VNS: vagus nerve stimulation; IV: Intravascular; LPS: Lipopolysaccharide; ChAT: Choline acyl transferase; Vglut2: Vesicular glutamate transporter 2; ChR: Channel rhodopsin; AR: Adrenergic receptor; TNF: Tumor necrosis factor alpha; NET: Norepinephrine transporter.

## Supplementary Methods

### Algorithm for determining the integration window for i°

The window for integrating i° over time is defined by 2 boundaries. The first (early) boundary is the onset of the stimulus train. The second (late) boundary is determined by algorithm operating over several steps. 1) determine if NE signal exists and if so, place a temporary estimated boundary, 2) determine oxidation potential (E°) of NE unique to the experimental setup, 3) calculate i° for each voltammogram within rough boundaries and place final boundary based on the maximum signal, 4) integrate over final boundaries to find Q°.

For the first step, the NE signal had to be characterized so that it could be detected automatically. A standard voltammogram for NE was defined by averaging the resulting voltammograms from 5 venous NE injection experiments. This standard voltammogram was normalized and the dot product between itself and each voltammogram in the time series of an experiment was used to determine the presence, or absence, of the signal. After the stimulus trigger, the first zero reached by the dot product trace was considered the temporary estimated boundary. If the dot product trace is never positive after the trigger stimulus, Q° defaults to zero. Each voltammogram between the temporary boundaries were averaged and used for the 2^nd^ step of the algorithm.

For the 2^nd^ step, MATLAB’s peak detection algorithm is applied to the average voltammogram from step 1, to determine the voltage corresponding to the peak (E°) and trough of the current curve generated by the oxidation of NE (points a and b in Figure 1b). To ensure that the selection of E° was not arbitrary, a mean and standard deviation was calculated from standard E° values from the 5 voltammograms used to define the standard voltammogram in step 1. In any experiment where E° was more than 3 standard deviations from the mean of this distribution, Q° defaults to zero.

For the 3^rd^ step, oxidation current (i°) is calculated for each voltammogram in the time series as the peak to trough difference in current. To determine the final integration boundary, the i° trace was smoothed with a moving average filter of size 10s and MATLAB’s peak detection algorithm is applied to find the time point of the first peak after the stimulus trigger. The final integration boundary is placed at the first time point where i° decreases to 10% of its first peak value after the first peak. If no time point satisfies this condition, Q° defaults to zero.

This algorithm was used to quantify data over the range of all data collected to minimize bias, however a human operator double checked all results and made manual corrections whenever appropriate.

Algorithm implemented in MATLAB with all of the raw data can be found here: https://github.com/mgerber000/SpleenFSCV

## Declarations

### Ethics approval and consent to participate

Animal experiments were conducted according to the US National Institutes of Health guidelines and were approved by the Institutional Animal Care and Use Committee (IACUC) of the Feinstein Institute for Medical Research (Protocol # 2019-010).

### Consent for publication

Not applicable

### Availability of data and materials

The analysis algorithms, along with all of the raw data can be found here: https://github.com/mgerber000/SpleenFSCV

### Competing interests

The authors declare that they have no competing interests.

## Funding

This research was conducted using departmental funds from the Feinstein Institutes for Medical Research.

## Authors’ contributions

ITM, MG conceived, designed, and performed experiments, analyzed data, interpreted data, and wrote the paper. NJ conceived and designed experiments. SPP designed experiments. TD, CS, VAP, YAA interpreted the data and wrote the paper. SZ conceived and designed experiments, analyzed data, interpreted data, wrote the paper. All authors approved the final version.

## Acknowledgments

The authors thank Kip Ludwig and James Trevanthan from University of Wisconsin-Madison for their helpful discussions, and Jason Wong from the Feinstein Institutes for assembling the optical cuff used in optogenetic stimulation experiments.

---

**Figure S1.**
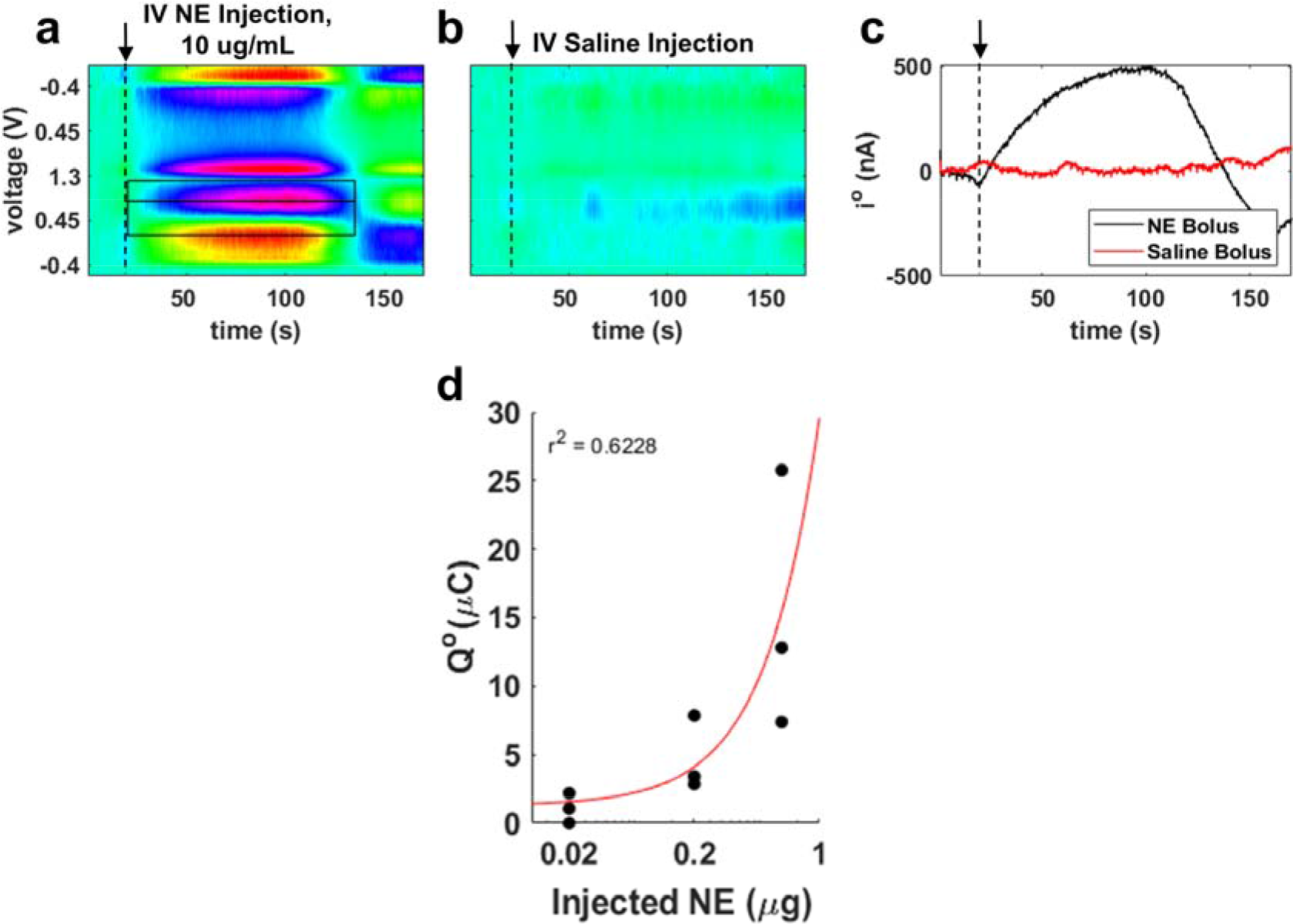
(a) Representative time-resolved voltammogram during intravenous bolus injection (10 μg/ml, 100 μL) of NE. (b) Time-resolved voltammogram during saline injection (100 μL). (c) Time course of i° in the experiments shown in b, c. (d) Amount of injected NE (μg) plotted against Q°.

**Figure S2.**
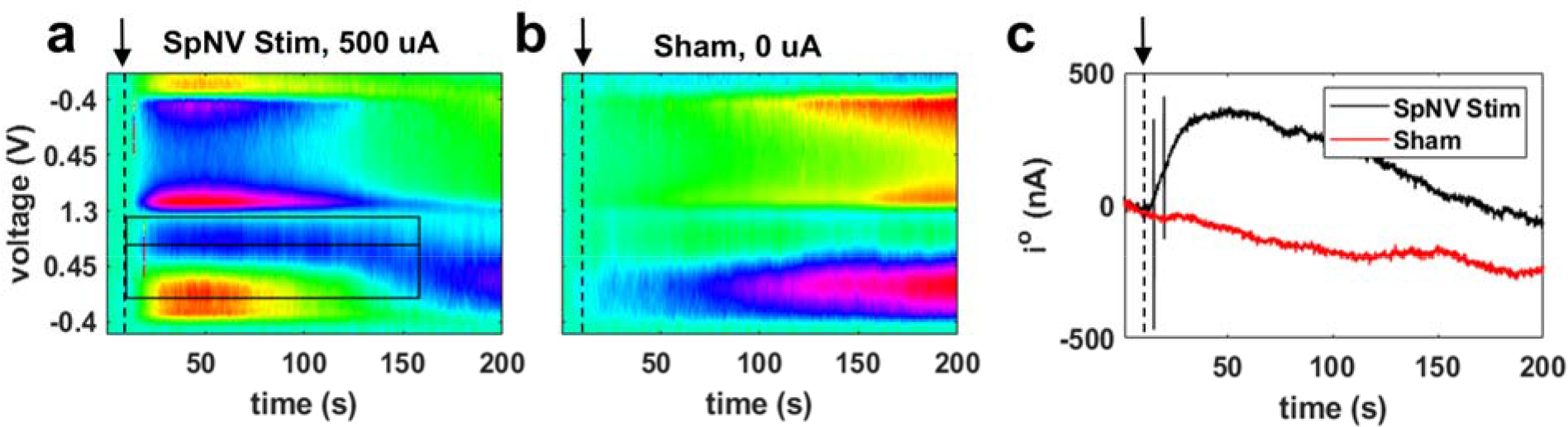
(a) Representative time-resolved voltammogram during SpNS (10 sec duration, 500 μs pulse width, 10 Hz pulsing frequency). (b) Time-resolved voltammogram during sham stimulation (recording of voltammogram without any stimulus applied). (c) Time course of i° in the experiments shown in a, b.

**Figure S3.**
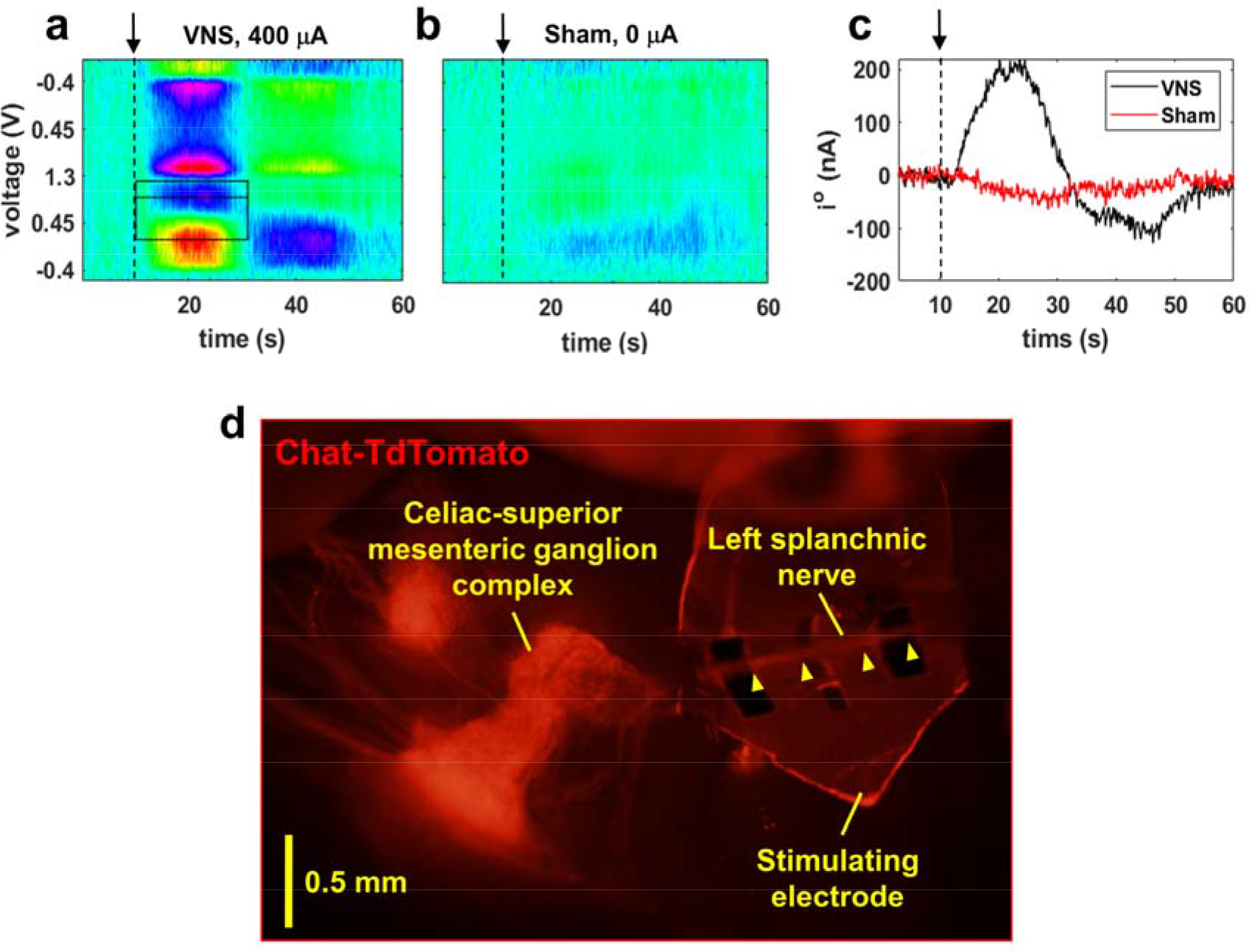
(a) The left celiac-superior mesenteric (CSM) ganglion complex is identified using fluorescence microscopy in ChAT-tdTomato mice (ChAT^+^ tissue appears in red). The splanchnic nerve is isolated and cuffed with a bipolar stimulating electrode. (a) Representative time-resolved voltammogram during left cervical VNS. (b) Time-resolved voltammogram during sham stimulation (recording of voltammogram without any stimulus applied). (c) Time course of i° in the experiments shown in b, c. Dashed vertical line indicates the onset of VNS. (d) The left celiac-superior mesenteric (CSM) ganglion complex is identified using fluorescence microscopy in ChAT-tdTomato mice (ChAT^+^ tissue appears in red). The splanchnic nerve is isolated and cuffed with a bipolar stimulating electrode.

## Notes

https://github.com/mgerber000/SpleenFSCV

### Competing Interest Statement

The authors have declared no competing interest.

### Summary of Updates

(1) Updated title. (2) Results and figures edited and improved. (3) Revised introduction and discussion. (4) New data added.

